# Peak-agnostic high-resolution cis-regulatory circuitry mapping using single cell multiome data

**DOI:** 10.1101/2023.06.23.544355

**Authors:** Zidong Zhang, Frederique Ruf-Zamojski, Michel Zamojski, Daniel J. Bernard, Xi Chen, Olga G. Troyanskaya, Stuart C. Sealfon

## Abstract

Single same cell RNAseq/ATACseq multiome data provide unparalleled potential to develop high resolution maps of the cell-type specific transcriptional regulatory circuitry underlying gene expression. We present CREMA, a framework that recovers the full cis-regulatory circuitry by modeling gene expression and chromatin activity in individual cells without peak-calling or cell type labeling constraints. We demonstrate that CREMA overcomes the limitations of existing methods that fail to identify about half of functional regulatory elements which are outside the called chromatin “peaks”. These circuit sites outside called peaks are shown to be important cell type specific functional regulatory loci, sufficient to distinguish individual cell types. Analysis of mouse pituitary data identifies a Gata2-circuit for the gonadotrope-enriched disease-associated Pcsk1 gene, which is experimentally validated by reduced gonadotrope expression in a gonadotrope conditional Gata2-knockout model. We present a web accessible human immune cell regulatory circuit resource, and provide CREMA as an R package.

---

Elucidating the mechanisms underlying the regulation of gene expression is fundamental for understanding the molecular basis of cell type identity, biological processes and disease. Cis-gene regulatory circuits, which consist of transcription factors (TFs) and their interactions with specific cis-regulatory sites on chromatin, serve a major role in determining gene expression ^1^. RNA-seq and ATAC-seq multiome technology, by profiling the regulatory circuit components within each nucleus, ^2,3^ sets the stage for reconstructing cell type-specific gene control circuitry at single cell resolution.

Analysis of single cell data typically initially reduces the search space by first calling chromatin peaks in pseudo-bulk data ^4–6^. Studies of ChIP-seq data have shown that weak binding sites, while functionally important, are often missed by genome-wide peak calling methods ^7–9^. We speculated that for single cell ATAC-seq data, the peak calling algorithms also may fail to identify many open or partly open regulatory loci that do not reach the statistical significance required for differential accessibility calling. Our evaluation of this possibility using functional domain databases indicated that restricting the circuit search to functional peaks neglects about half of known functional regulatory regions. Accordingly, a framework that does not require peak calling is desirable to leverage the power of single cell multiome datasets for understanding gene control mechanisms.

To address this bottleneck, we developed CREMA (Control of Regulation Extracted from Multiomics Assays), a framework for the systematic survey of gene regulatory circuits from single cell multiomics data. CREMA recovers circuitry by modeling gene expression and chromatin accessibility directly over the entire cis-regulatory region, without being restricted by either peak calling or cell type identification. We demonstrate the improvement of regulatory circuit recovery by CREMA relative to the current state-of-the-art method and show the value of CREMA for identifying new circuitry and accessibility site variation that defines individual cell types. Applying CREMA to mouse pituitary data, we show how it can identify cell type specific circuits and identify a gata2-circuit regulating a disease-associated target that is validated in a conditional gata2 mouse knockout model. In addition to making CREMA available to the research community (https://github.com/zidongzh/CREMA), we use CREMA to generate a web-accessible research resource comprising the regulatory circuitry of human blood immune cells (https://rstudio-connect.hpc.mssm.edu/crema-browser/).

## Results

### Motivation

Each gene regulatory circuit consists of a TF, a cis-regulatory domain that interacts with the TF, and a target gene that has altered transcription resulting from this interaction. Multiple circuits involving the same TF binding at different locations or multiple TFs interacting at the same or different loci are the major cis-regulatory mechanisms regulating gene expression. Existing gene control circuit analysis methods only identify the potential regulatory domains for these circuits that are contained within called chromatin peaks in ATAC-seq data. In order to assess the degree to which this restriction may limit identification of cis-regulatory domains and their associated circuits, we investigated the fraction of known regulatory loci in human blood that are outside of called chromatin peaks. We determined the proportion of known functional domains in two reference databases that were contained within called chromatin peaks using high resolution reference single cell ATACseq data (see Online Methods). A majority of eQTLs in the GTEX DAPG fine-mapped eQTL database ^10^ and of enhancers in the EnhancerAtlas ^11^ database are located outside of the peaks called using reference high resolution human peripheral blood mononuclear cell (PBMC) chromatin accessibility data ^12,13^ (Fig. 1A). We observed similar results in other fine-mapped eQTL and enhancer databases (Supplementary Fig. 1). These results suggest that multiome circuit inference methods that rely on chromatin peak calling will miss about half of the regulatory landscape and circuitry underlying gene control. To address this gap, we developed CREMA to improve the reconstruction of gene regulatory circuitry.

**Figure 1.**
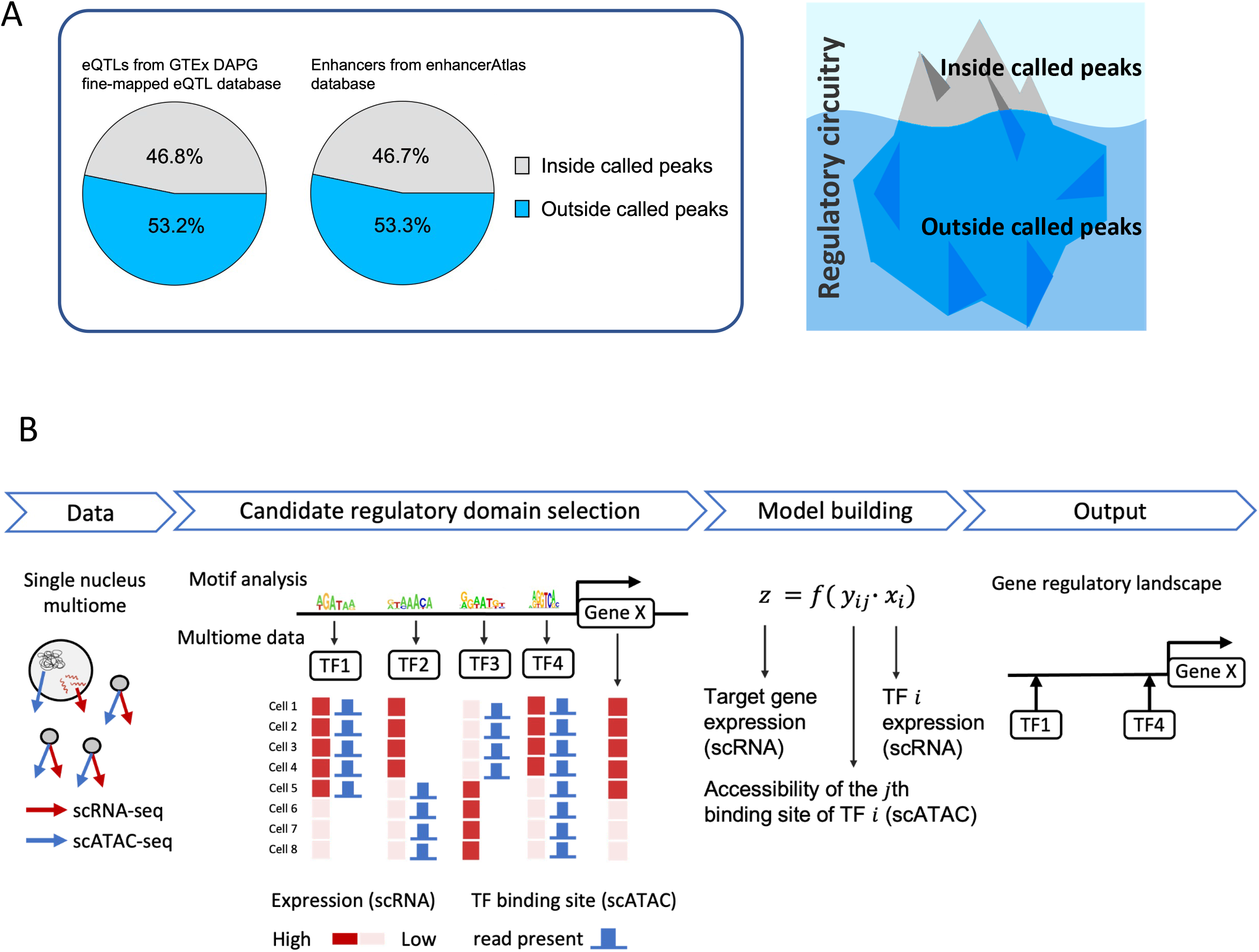
Motivation and workflow of CREMA. A: Percentage of eQTLs and enhancers from gold standard databases located inside and outside of ATAC peaks called in a human PBMC single nucleus multiome data. Reference blood eQTLs are obtained from the GTEx DAPG fine-mapped eQTLs database. Reference blood enhancers are obtained from the enhancerAtlas database. B: Schematic of the CREMA method. CREMA takes single nucleus multiome (RNAseq + ATACseq within each cell) as input. It scans for potential cis-TF binding sites by motif analysis. It then fits a linear model for gene expression as a function of chromatin accessibility and TF expression to each cell in the dataset to select highly significant regulatory circuits. The circuits identified are supported by the coincidence of TF expression, binding site accessibility and target gene expression within individual cells.

### CREMA Framework

CREMA was designed to identify transcriptional regulatory circuits over the entire cis-regulatory region of each gene. CREMA finds circuits that are supported by the co-incidence of TF expression, target gene expression and binding site accessibility in individual cells. A schematic of the method is shown in Fig. 1B. CREMA first selects the target genes to model that have detectable expression above a threshold in a minimum number of cells and proportion of all cells (See Methods). For each of these target genes, CREMA uses motif analysis to select potential TF binding sites in a +/- 100kb window surrounding the transcription start site (TSS).

Each site, together with the TF and gene constitute a potential regulatory circuit. A linear model for each potential circuit is constructed where the expression of each gene in each cell is a function of the expression of the TF and the binarized accessibility in a 400 bp window centered on the site. Using all the cells in the dataset, the TF-site-gene circuits showing the best fits are selected (See Methods).

### Benchmarking

Because CREMA does not rely on a predefined set of chromatin peaks called at the pseudo-bulk level, it has the potential to recover many more regulatory domains compared to analyses relying on peak calling. We used CREMA to analyze single cell blood multiome data and found regulatory circuits both inside and outside of chromatin peaks. The number of circuits identified within peaks was comparable to that obtained using the currently available multiome regulatory circuit discovery method, which relies on peak calling ^12^. CREMA also identified a large number of circuits that are outside of called peaks, which cannot be found with a peak-calling dependent method(Fig. 2A).

**Figure 2.**
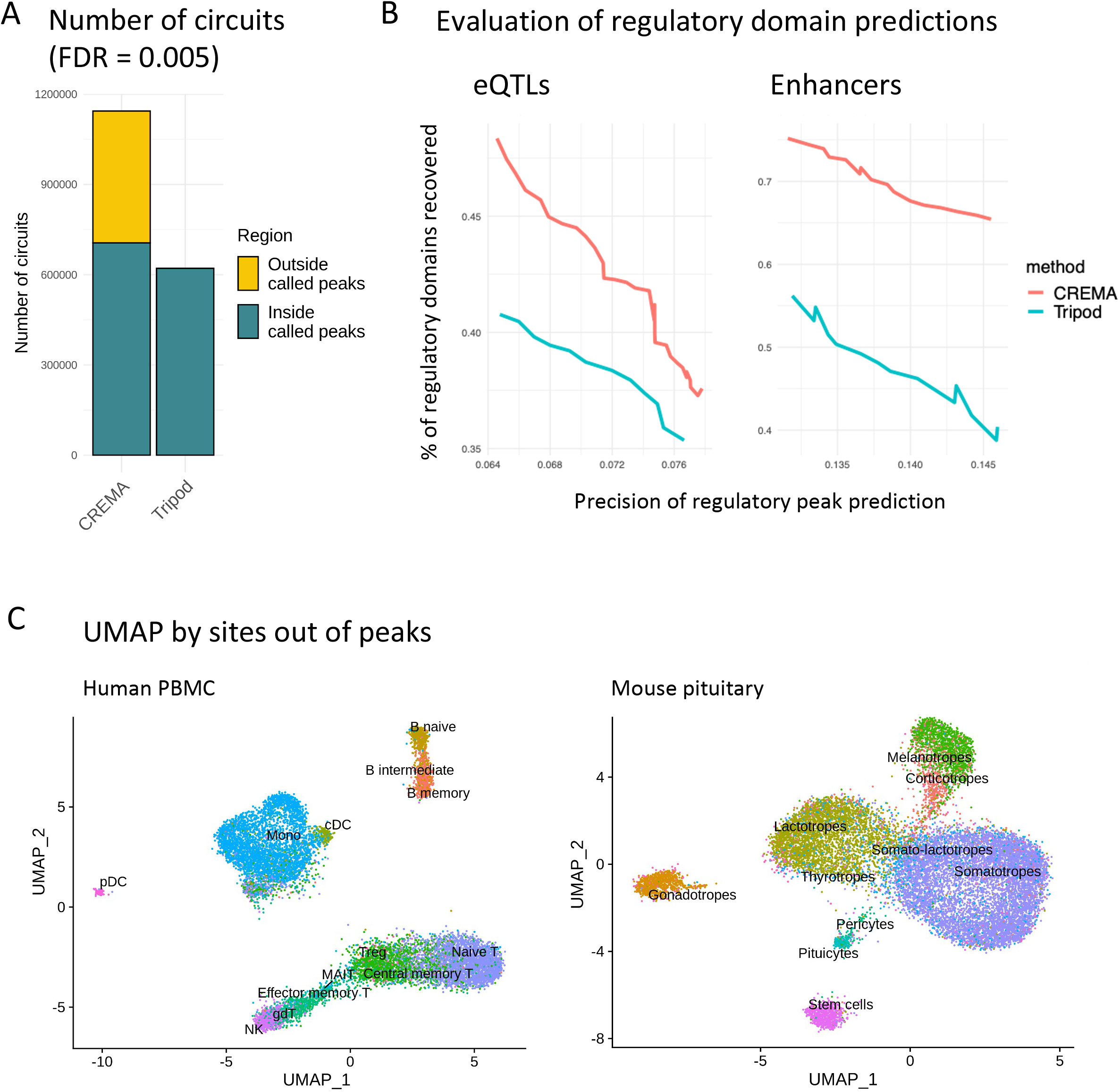
CREMA performance and utility. A: Number of regulatory circuits identified by TRIPOD 12 and CREMA at FDR cutoff = 0.005. The circuits from CREMA were categorized as “inside called peaks” or “outside called peaks” depending on whether the binding site of the circuit overlapped with any chromatin peak. Because the circuit inference from TRIPOD was restricted to the chromatin peaks, all the circuits from TRIPOD are inside called peaks. B: Percentage of true regulatory regions recovered by TRIPOD and CREMA when controlling for the precision in the peak regions. Predictions from the two methods were selected at different FDR cutoffs to calculate the precision of regulatory peak prediction and recovery of true regulatory regions from the reference gold standards (see methods). Reference blood eQTLs are obtained from the GTEx DAPG fine-mapped eQTLs database. Reference blood enhancers are obtained from the enhancerAtlas database. C: Cis-regulatory domains outside of called peaks resolve major cell types in human PBMC and mouse pituitary. UMAP dimension reductions were calculated by using only the accessibilities of CREMA identified cis-regulatory domains outside of ATAC peaks as features. Cell type annotations were from independent analysis using the expression of known marker genes (see methods).

The importance of the additional regulatory landscape recovered by CREMA was evaluated using gold standard functional domain databases. CREMA greatly improved recovery of circuits acting at functional domains in both reference eQTL and enhancer databases (Fig. 2B, Supplementary Fig. 2). To further assess the importance of the extra-peak regulatory circuitry that CREMA recovers, we evaluated whether the regulatory circuit chromatin domains identified by CREMA that were outside of called chromatin peaks contributed to cell type specification. In addition to the PBMC dataset analysis shown in Fig. 2A, we generated a mouse pituitary multiome dataset that was also analyzed using CREMA. In both cases, we used only the chromatin regulatory sites discovered by CREMA that are outside of called peaks as features for UMAP projections. We found that in both tissues, the major cell types were distinguishable (Fig. 2C). These results indicate that the comprehensive circuitry mapping achievable with CREMA is necessary to elucidate the gene control mechanisms underlying the differences among cell types.

### CREMA identified Gata2 circuit

We next investigated the regulatory circuits identified by CREMA in pituitary involving the pioneer TF, Gata2 ^14^. In pituitary, Gata2 is necessary for gonadotrope lineage specification and regulates the production of follicle-stimulating hormone. In mouse pituitary single cell multiome data, CREMA identified circuits regulating 323 target genes. Because Gata2 was highly expressed in both gonadotrope and somatrope cell types (Supplementary Fig. 3), we focused on the circuits in these two cell types for validation. Among the 323 target genes in Gata2 circuits, 88 were highly expressed in the gonadotropes and 200 were highly expressed in the somatotropes (Fig. 3A). To validate these CREMA predicted circuits, we assessed the expression of the target genes for these circuits in single cell RNAseq data obtained from a gonadotrope-specific conditional Gata2 knockout ^15^. In this knockout, Gata2 function was absent in gonadotropes, and 10 of the predicted gonadotrope Gata2 target genes were significantly down regulated. In contrast, Gata2 function was preserved in somatotropes and none of the predicted Gata2 target genes showed significant down regulation (p = 3.5×10^−6^, z-test of two proportions, Fig. 3A). These results provide strong support for the recovery of the Gata2 circuitry by CREMA.

**Figure 3.**
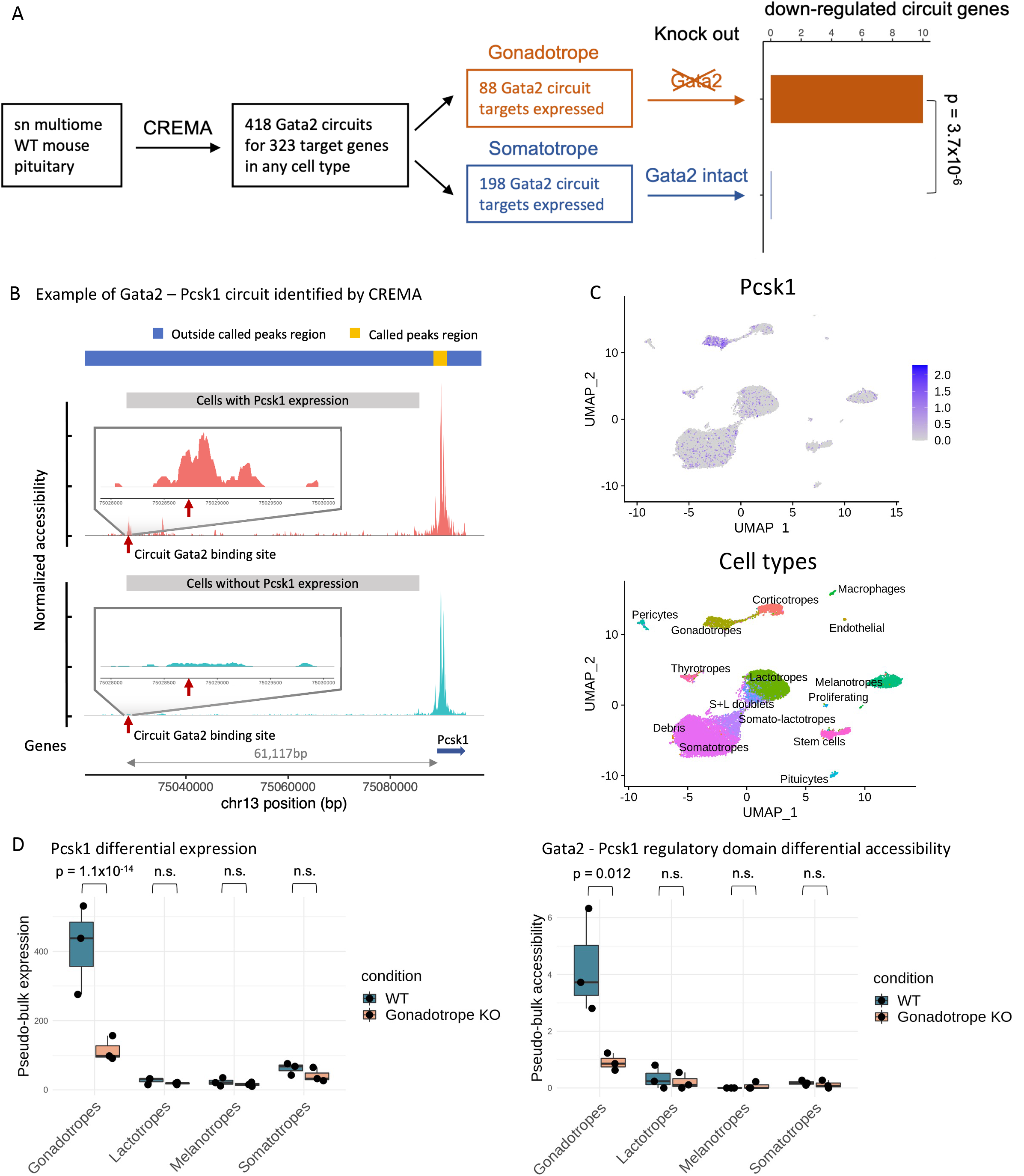
Gata2 - Pcsk1 circuit in the pituitary gonadotrope cells. A: Schematic showing the analysis of Gata2 circuits by CREMA in the mouse pituitary and validation by differentially expressed genes in the conditional Gata2 knockout data. (p = 3.5×10-6, Z = 4.5, df = 1, one-sided z-test of two proportions) B: Detailed view of a CREMA identified Gata2-Pcsk1 circuit where Gata2 interacts with a cis-regulatory domain located ∼61kb upstream of the TSS of Pcsk1. Normalized accessibilities were plotted separately for cells with and without Pcsk1 expression. Zoomed in plot showing the detailed chromatin accessibility pattern around the Gata2 binding site (red arrow). C: UMAPs showing the expression of Pcsk1 in the pituitary cells and the cell type annotations. D: Box plot and point plot showing the pseudobulk RNA of Pcsk1 and pseudobulk ATAC of the Gata2 site in each cell type of the wild type mouse pituitary samples (n = 3) and gonadotrope-conditional Gata2 knockout samples (n = 3).

We next focused on the Gata2 circuit involved in the regulation of the Pcsk1 gene, which is implicated in infertility, obesity and diabetes ^16–18^. CREMA identified a significant cis-regulatory domain with a Gata2 binding motif at 61kb upstream of the Pcsk1 TSS. This domain was highly accessible in cells with Pcsk1 expression but was not included within called peak regions and could not have been identified by a peak-calling dependent method (Fig. 3A). Pcsk1 was expressed in multiple cell types in the pituitary: gonadotropes, lactotropes, melanotropes and somatotropes (Fig. 3B). However, the expression of Pcsk1 and the accessibility of this cis-regulatory domain were down regulated only in the gonadotropes in the conditional knockout data, where Gata2 activity was eliminated, while remaining unchanged in the other cell types (Fig. 3C). These results demonstrate the usefulness of CREMA for leveraging single cell multiome data to obtain insight into the regulatory circuitry controlling gene expression at cell type resolution.

### Regulatory circuitry resource for human immune cells

The orchestration of the immune response in health and disease depends on the modulation of gene expression in the different immune cell types. In order to provide a resource for the study of gene regulatory mechanisms in immune cells, we used CREMA to identify the regulatory circuitry in blood using a single cell multiome dataset and provide this analysis as a community resource. Circuitry can be summarized both in a TF-centric and gene-centric manner. We first summarized the CREMA regulatory circuits in a TF-centric perspective, defining a TF module as the collection of regulatory circuits sharing a common TF in each cell type (see Methods). Selected TF modules and their activities in the major immune cell types are presented in Figure 4A.

**Figure 4.**
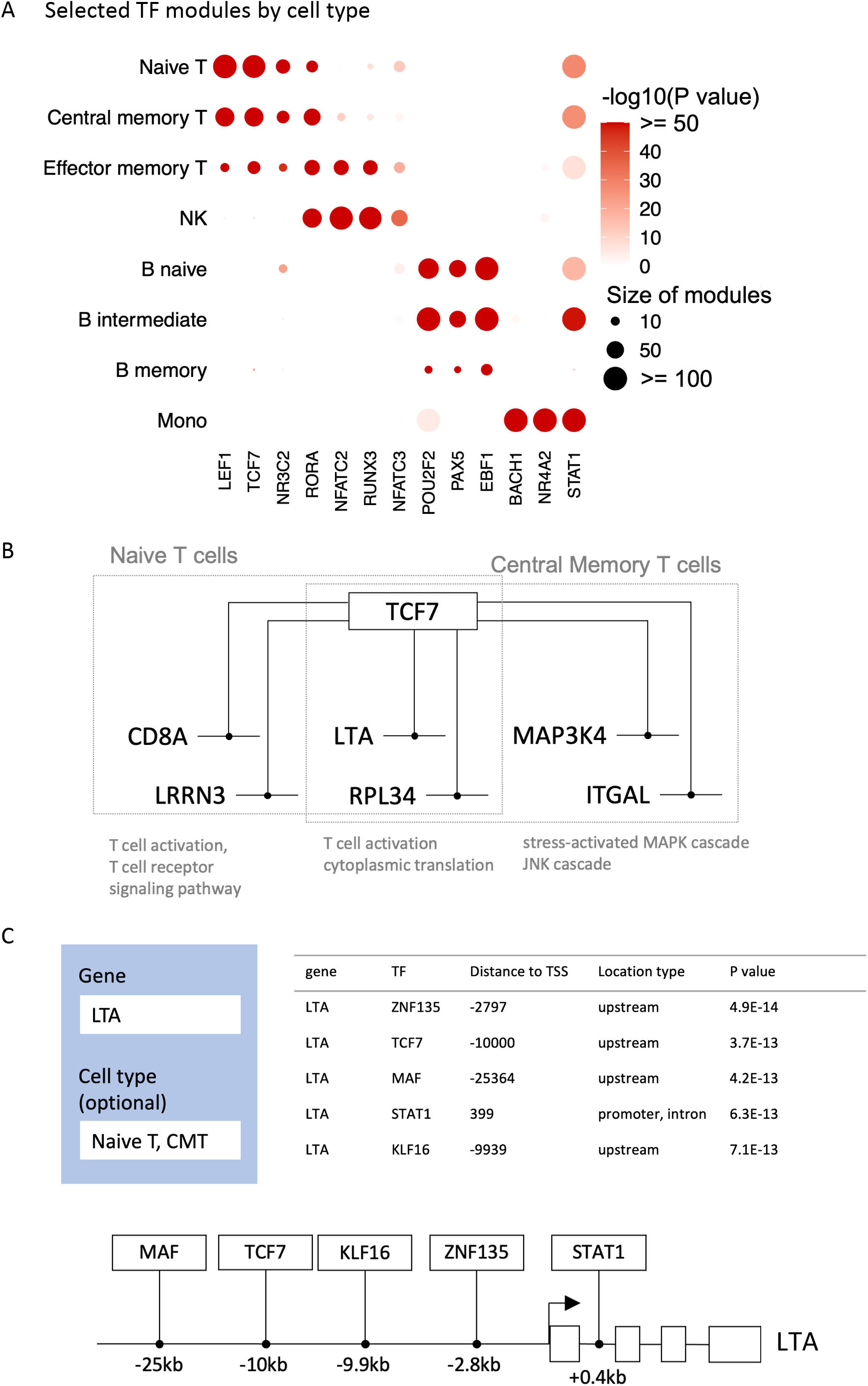
Regulatory circuitry of human immune cells. A: Selected CREMA identified TF modules and their activities in immune cell types. B: Selected CREMA identified regulatory circuits in the TCF7 module that are shared between naive T cells and central memory T cells, and circuits in the TCF7 module that are specific to one of the two cell types. GO terms annotated to these target genes are labeled below. C: Example of a queried gene LTA and the list of CREMA identified regulatory circuits targeting this gene.

As an example, we focused on the TCF7 module, which is active mainly in the naive T cells and central memory T cells. Within the TCF7 module, there were circuits shared by the two cell types, such as the circuit regulating the target gene LTA which encodes a cytokine expressed by resting and activated T cells ^19,20^ (Fig. 4B). There were also TCF7 circuits specifically active in one of the cell types. For example, the TCF7-CD8A circuit was active only in the naive T cells and CD8A is involved in T cell activation. The TCF7-MAP3K4 circuit was active only in the central memory T cells and MAP3K4 is involved in the stress-response MAPK cascade (Fig. 4B, see Supplementary Table 1).

A full picture of the gene control within each cell type is obtained by aggregating the multiple regulatory circuits involved in the control of specific genes. In the immune cell resource, we provide access to the entire regulatory circuitry within each cell type. The user can query a gene of interest to obtain a list of regulatory circuits targeting this gene, including the TF and the locations of the cis-regulatory domains interacting with these TFs. We show an example of a query gene LTA and the top five regulatory circuits identified by CREMA (Fig. 4C). This immune cell resource is designed to help the research community generate hypotheses about the gene control mechanisms specific to immune cell subtypes and may also help the selection of specific TFs to target for therapeutic immune modulation.

## Discussion

CREMA leverages single cell multiome data to infer cis-regulatory circuitry covering the entire cis-regulatory region. CREMA identifies cis-regulatory domains by directly combining the local chromatin accessibility of potential TF binding sites and TF expression without relying on calling ATAC peaks. This expanded search space enables the identification of the large proportion of regulatory circuitry outside of called peaks that contributes to gene control and to cell type specification. The performance of CREMA has been validated using public functional domain databases and a conditional knockout model and an immune cell gene circuitry analysis has been developed as a public resource.

For circuits that are located within peaks, because CREMA models local chromatin accessibility of the TF binding site in a small chromatin window relative to the peak region, CREMA provides higher resolution of the chromatin domain for the circuit than peak-calling dependent approaches. In addition to being chromatin peak-agnostic, the CREMA framework is cell type agnostic. Cell type identification is utilized only after performing the CREMA analysis in order to evaluate the cell type specificity of the circuits identified. This gives CREMA the potential to identify circuits in poorly represented or unlabeled cell types or unlabeled cell types.

We have developed a resource of the full regulatory circuitry of human immune cells to facilitate hypothesis generation and experiment design for the immune research community (https://rstudio-connect.hpc.mssm.edu/crema-browser/). CREMA, publicly available via an R package (https://github.com/zidongzh/CREMA), can help realize the potential of multiome datasets to resolve the circuitry underlying gene control in individual cells.

## Competing Interests

S.C.S. is a founder of GNOMX Corp and serves as chief scientific officer. The remaining authors declare no competing interests.

## Acknowledgments

We thank the Single-cell and Spatial Technologies team at the Center for Advanced Genomics Technology, Department of Genetics and Genomic Sciences, the Icahn School of Medicine at Mount Sinai for providing the experimental, computational, data resources, and staff expertise. This work is supported by the Defense Advanced Research Projects Agency under contract N6600119C4022 (S.C.S. and O.G.T.) NIH grant RO1 DK46943 (S.C.S) and NIH grant R01GM071966 (O.G.T) and Simons Foundation grant 395506 (O.G.T).

## Online Methods

### CREMA framework

#### Gene filtering

We focused on modeling genes and TFs above a certain level of expression in the dataset. Specifically, we applied 2 filters on the genes: 1) the gene counts must be non-zero in at least 0.1% of the cells or 3 cells, whichever was larger, and 2) the gene total count in all cells should be larger than (0.2% × total cell number).

#### Candidate regulatory domain selection

For each target gene, we analyzed the entire +/- 100kb window around the transcription start site (TSS) without reference to ATAC-seq peak calling. We scanned for potential TF binding sites in this region by motif analysis. Specifically we used the human TF position weight matrices from the JASPAR database and mouse TF position weight matrices from the CIS-BP database. For the motif analysis we used the function matchMotifs from the r package motifmatchr having p<5e-5.

#### Model building

To select regulatory circuits supported by the co-incidence of TF expression, target gene expression and binding site accessibility, we used a linear regression framework where the level of TF is weighted by the accessibility of that TF’s binding site. Specifically, for each TF and each binding site found in the candidate regulatory domains, we counted the number of ATACseq cut sites overlapping with a 400bp window centered around the binding site in each single cell, and binarized the results as open (counts >= 1) or closed (counts = 0). Then the level of TF RNA and the accessibility of TF binding sites were combined in a linear regression:

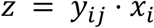

Where z is the RNA level of the target gene, *x*_*i*_ is the RNA level of the *i*th TF, and *y*_*ij*_ is the binarized chromatin openness of the *j*th binding site of the *i*th TF in the candidate regulatory regions. The RNA levels used in the model are normalized RNA levels with SCTranscform. The rationale was that TFs with a closed binding site would not be selected as significant regulators in this framework. Because many TFs had more than one binding site, there was high collinearity among the regressors. Therefore we evaluated the significance of each TF-site combination by linear regression individually and reported all significant TF-site combinations, instead of using a multi-regression framework.

### Data and preprocessing

#### Human PBMC data from 10X Genomics

The single nucleus multi-omics dataset of human PBMC was provided by 10X Genomics as a reference dataset. Specifically, the dataset “pbmc_granulocyte_sorted_10k” processed using CellRanger v1.0.0 was downloaded from 10X Genomics, and it was processed following the vignette “Joint RNA and ATAC analysis: 10x multiomic” from the r package Signac v1.5.0.

#### Single nucleus multiome (RNA+ATAC) of male mouse pituitary

The pituitary used in this study was collected from a male C57BL/6 mice aged 10 weeks. Animals were on a 12-hour on, 12-hour off light cycle (lights on at 7 AM; off at 7 PM). Once collected, the pituitary was immediately snap-frozen following dissection, and stored at -80C until the assay was started.

Nuclei isolation was performed as described in ^21,22^. Briefly, the snap-frozen pituitary was thawed on ice. RNAse inhibitor (NEB MO314L) was added to the homogenization buffer (0.32 M sucrose, 1 mM EDTA, 10 mM Tris-HCl, pH 7.4, 5mM CaCl2, 3mM Mg(Ac)2, 0.1% IGEPAL CA-630), 50% OptiPrep (Stock is 60% Media from Sigma; cat# D1556), 35% OptiPrep and 30% OptiPrep right before isolation. The pituitary was homogenized in a dounce glass homogenizer (1ml, VWR cat# 71000-514), and the homogenate filtered through a 40 m cell strainer. An equal volume of 50% OptiPrep was added, and the gradient centrifuged (SW41 rotor at 9200rpm; 4C; 25min). Nuclei were collected from the interphase, washed, resuspended in 1X nuclei dilution buffer (10X Genomics), and counted (Nexcelom K2 counter).

Sn multiome was performed following the Chromium Single Cell Multiome ATAC and Gene Expression Reagent Kits V1 User Guide (10x Genomics, Pleasanton, CA) on a male mouse wild-type sample. Nuclei were counted as described above, transposition was performed in 10 l at 37C for 60min targeting 10,000 nuclei, before loading of the Chromium Chip J (PN-2000264) for GEM generation and barcoding. Following post-GEM cleanup, the library was pre-amplified by PCR, after which the sample was split into three parts: one part for generating the snRNAseq library, one part for the snATACseq library, and the rest was kept at -20C. SnATAC and snRNA libraries were indexed for multiplexing (Chromium i7 Sample Index N, Set A kit PN-3000262, and Chromium i7 Sample Index TT, Set A kit PN-3000431 respectively). The library was quantified by Qubit 3 fluorometer (Invitrogen) and quality was assessed by Bioanalyzer (Agilent). This library was sequenced first in a Miseq (Illumina) to assess the reads and balance the sequencing pool, then it was sequenced in a Novaseq 6000 (Illumina) at the New York Genome Center (NYGC) following 10X Genomics recommendations.

The sequencing data was preprocessed with cellranger-arc-2.0.0. The dataset was then processed as described by the vignette “Joint RNA and ATAC analysis: 10x multiomic” from the r package Signac v1.5.0. Cell types were identified by label transfer from a well annotated single nucleus RNAseq dataset^21^ using the r package Seurat v4.1.0

#### Single nucleus RNAseq and ATACseq of WT and Gata2KO mice

Processed single nucleus RNAseq and single nucleus ATACseq datasets of 3 wild type mice (WT) and 3 mice with Gata2 conditionally knocked out in the gonadotrope cells of the pituitary were provided by Daniel Bernard’s lab at McGill University^15^. Cell clusters corresponding to the gonadotropes were located using marker genes of gonadotropes as described before.

### Benchmarking

#### Number of discoveries

We ran both CREMA and TRIPOD on a human PBMC sn multiome dataset. We use the same FDR cutoff of 0.005 on both methods. For TRIPOD, we selected all the TF-peak-gene combinations passing the FDR cutoff and each of these combinations was counted as one regulatory circuit. For CREMA, we selected all the TF-site-gene combinations passing the FDR cutoff, and overlaid the site location to chromatin peaks to determine where the regulatory circuit is within peak regions or outside of peak regions.

#### Public databases of true regulatory regions

EnhancerAtlas was downloaded from EnhancerAtlas 2.0 database and all the enhancer-gene interactions in blood cell types were combined. Fantom and 4D genome databases were downloaded from the processed datasets provided by the TRIPOD package. Fine-mapped eQTLs were downloaded from GTEx v8. See supplementary table 2 for the URLs of these databases.

#### Recovery of true regulatory regions

We applied CREMA and TRIPOD to the human PBMC sn multiome dataset to extract regulatory regions for the top 1000 variable genes. Specifically we ran TRIPOD with default settings and extracted all regulatory peaks with both level 1 and level 2 testings. Three enhancer databases and three fine-mapped eQTL databases described in the last section were used to evaluate the precision of regulatory region predictions and recovery of the true regulatory regions. To compare across the two methods, we evaluated the performance from the two methods by setting different FDR cutoffs in the range of 0.1 to 0.0001. For each FDR cutoff, we calculated: 1) the recovery of true regulatory regions, defined as the percentage of true regulatory regions from the databases that overlap with the regulatory regions predicted by TRIPOD and CREMA. 2) precision of predictions, defined as the percentage of predicted regions that overlap with true regulatory regions from the databases.

Specifically, chromatin peaks predicted by TRIPOD are larger in sizes than the regulatory sites predicted by CREMA, and larger regions are more likely to overlap with a true regulatory region from the reference databases. So to make the calculation of the precision of prediction in the same space for TRIPOD and CREMA, we converted the regulatory sites predicted by CREMA to the chromatin peaks that overlapped with these sites for calculating the precision of predictions. If a chromatin peak overlapped with multiple sites from CREMA, we used the minimum p value among these sites as the p value for this peak.

#### Recovery of cell types

We applied CREMA on the human PBMC sn multiome dataset and the mouse pituitary sn multiome dataset. In both cases, we extracted regulatory circuits with an FDR cutoff of 0.0001 and selected cis-regulatory regions outside of the called chromatin peaks. We then calculated the chromatin accessibility in these regions and used them as features for LSI and UMAP dimension reduction on the datasets. In the UMAP visualization, the cells were colored by the original cell type annotations obtained by label transfer from reference datasets as described in the “Data and preprocessing” section.

### Gata2 regulatory circuits in the pituitary

#### Extraction of Gata2 circuits in the pituitary cell types

We applied CREMA to the sn multiome dataset of wildtype mouse pituitary. We extracted all the regulatory circuits with an FDR cutoff of 0.0001. We selected all the regulatory circuits where Gata2 was the TF. In this dataset, there were 866 gonadotrope cells and 7420 somatotrope cells. For gonadotropes, we determined target genes of Gata2 as active in gonadotropes if they were detected in at least 260 cells (30%) of the gonadotropes. We used the same cutoff of 260 cells to determine Gata2 targets as active in the somatotropes. We chose to use the number of cells detected as the cutoff in order to accommodate possible higher heterogeneity within the somatotrope cells. The cell type specific target genes were analyzed for differential expression between the wild type and conditional knockout datasets.

#### Differential analysis of the Gata2-Pcsk1 circuit

We compared the expression of Pcsk1 and the accessibility of the Gata2 cis-regulatory site chr13:75028714-75028724 between the 3 wild type samples and 3 conditional knockout samples by pseudobulk analysis. The single cell expression and accessibilities were summed at cell type resolution and differential analysis were performed using DESeq2.

### Regulatory circuitry of the human immune cells

#### Regulatory circuits in PBMC

We applied CREMA to the sn multiome dataset of human PBMC. We selected regulatory circuits with a FDR cutoff of 0.0001.

#### Circuit activities and TF module activities in cell types

For visualizing the highly active TF modules in the major immune cell types, we calculated the circuit activities and TF modules activities in each cell type. The activity of each regulatory circuit in each cell was calculated by taking the product of the expression level of the TF, the expression level of the target gene and the binarized accessibility of the cis-regulatory site in the cell. To summarize the activities of regulatory circuits at cell type resolution, we used two methods: 1) a binary activity score where a regulatory circuit was defined as active in a cell type if it was active in more than 10% of the cells in that cell type and more than 50 cells of that cell type, 2) a continuous activity score where the activity of a regulatory circuit in a cell type was defined as the proportion of cells in that cell type where the regulatory circuit was active. To summarize the activities of regulatory circuits in a TF-centric view, we defined a TF module as the collection of all the regulatory circuits involving that TF. For each TF module and each cell type, we calculated 1) the number of active regulatory circuits in that cell type as measured by the binary activity score under that TF module, 2) the specificity of the regulatory circuits of that TF module to that cell type, measured by summing the continuous activity scores of the regulatory circuits and converting to a z score.

### Data and code availability

The lab generated single nucleus multiome dataset of mouse pituitary is accessible at GSE234943. CREMA is available as an R package at https://github.com/zidongzh/CREMA. The web-accessible resource of the regulatory circuitry of human blood immune cells is available at https://rstudio-connect.hpc.mssm.edu/crema-browser/. The source code for the analysis in this manuscript is available at https://github.com/zidongzh/CREMA_manuscript.

## References

1. Kim, H. D., Shay, T., O’Shea, E. K. & Regev, A. Transcriptional Regulatory Circuits: Predicting Numbers from Alphabets. Science 325, 429–432 (2009).

2. Ma, S. et al.. Chromatin Potential Identified by Shared Single-Cell Profiling of RNA and Chromatin. Cell 183, 1103–1116.e20 (2020).

3. Chen, S., Lake, B. B. & Zhang, K. High-throughput sequencing of the transcriptome and chromatin accessibility in the same cell. Nat. Biotechnol. 37, 1452–1457 (2019).

4. Stuart, T., Srivastava, A., Madad, S., Lareau, C. A. & Satija, R. Single-cell chromatin state analysis with Signac. Nat. Methods 18, 1333–1341 (2021).

5. Schep, A. N., Wu, B., Buenrostro, J. D. & Greenleaf, W. J. ChromVAR: Inferring transcription-factor-associated accessibility from single-cell epigenomic data. Nat. Methods 14, 975–978 (2017).

6. Bravo González-Blas, C. et al. cisTopic: cis-regulatory topic modeling on single-cell ATAC-seq data. Nat. Methods 16, 397–400 (2019).

7. Nakato, R. & Shirahige, K. Recent advances in ChIP-seq analysis: from quality management to whole-genome annotation. Brief. Bioinform. bbw023 (2016) doi:10.1093/bib/bbw023.

8. Landt, S. G. et al. ChIP-seq guidelines and practices of the ENCODE and modENCODE consortia. Genome Res. 22, 1813–1831 (2012).

9. Schmidt, D. et al. A CTCF-independent role for cohesin in tissue-specific transcription. Genome Res. 20, 578–588 (2010).

10. Wen, X., Pique-Regi, R. & Luca, F. Integrating molecular QTL data into genome-wide genetic association analysis: Probabilistic assessment of enrichment and colocalization. PLOS Genet. 13, e1006646 (2017).

11. EnhancerAtlas 2.0: an updated resource with enhancer annotation in 586 tissue/cell types across nine species | Nucleic Acids Research | Oxford Academic. https://academic.oup.com/nar/article/48/D1/D58/5628925.

12. Jiang, Y. et al. Nonparametric single-cell multiomic characterization of trio relationships between transcription factors, target genes, and cis-regulatory regions. Cell Syst. 13, 737-751.e4 (2022).

13. Hao, Y. et al. Integrated analysis of multimodal single-cell data. Cell 184, 3573-3587.e29 (2021).

14. Wu, D. et al. NAR Breakthrough Article: Three-tiered role of the pioneer factor GATA2 in promoting androgen-dependent gene expression in prostate cancer. Nucleic Acids Res. 42, 3607 (2014).

15. Schang, G. et al. Transcription factor GATA2 may potentiate follicle-stimulating hormone production in mice via induction of the BMP antagonist gremlin in gonadotrope cells. J. Biol. Chem. 298, (2022).

16. Folon, L. et al. Contribution of heterozygous PCSK1 variants to obesity and implications for precision medicine: a case-control study. Lancet Diabetes Endocrinol. 11, 182–190 (2023).

17. Severe obesity and diabetes insipidus in a patient with PCSK1 deficiency - ScienceDirect. https://www.sciencedirect.com/science/article/pii/S1096719213001145?via%3Dihub.

18. Genetic Variants in PCSK1 Gene Are Associated with the Risk of Coronary Artery Disease in Type 2 Diabetes in a Chinese Han Population: A Case Control Study | PLOS ONE. https://journals.plos.org/plosone/article?id=10.1371/journal.pone.0087168.

19. Ware, C. F., Crowe, P. D., Grayson, M. H., Androlewicz, M. J. & Browning, J. L. Expression of surface lymphotoxin and tumor necrosis factor on activated T, B, and natural killer cells. J. Immunol. Baltim. Md 1950 149, 3881–3888 (1992).

20. Ohshima, Y. et al. Naive human CD4+ T cells are a major source of lymphotoxin alpha. J. Immunol. Baltim. Md 1950 162, 3790–3794 (1999).

21. Ruf-Zamojski, F. et al. Single nucleus multi-omics regulatory landscape of the murine pituitary. Nat. Commun. 12, 2677 (2021).

22. Mendelev, N. et al. Multi-omics profiling of single nuclei from frozen archived postmortem human pituitary tissue. STAR Protoc. 3, 101446 (2022).

