## Supplementary figures and tables for "Peak-agnostic high-resolution cis-regulatory circuitry mapping using single cell multiome data"

Supp fig 1. Percentage of eQTLs and enhancers from gold standard databases that locate inside and outside of ATAC peaks called in a human PBMC single nucleus multiome data.

#### eQTLs

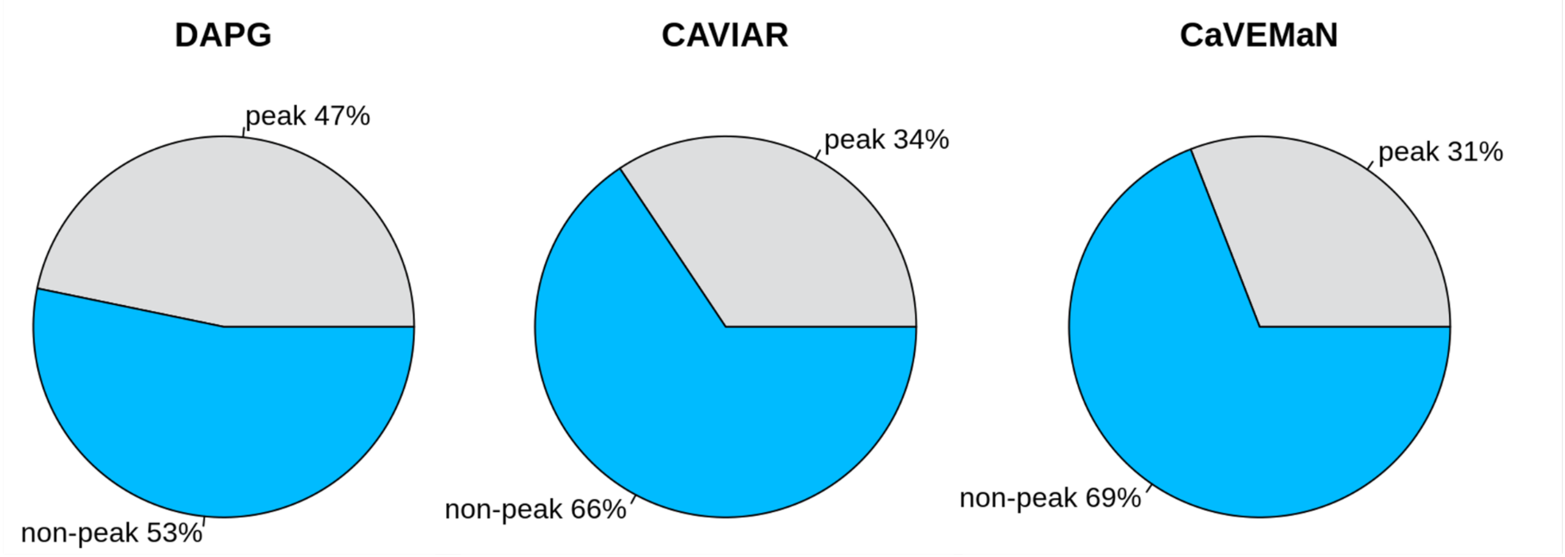

#### Enhancers

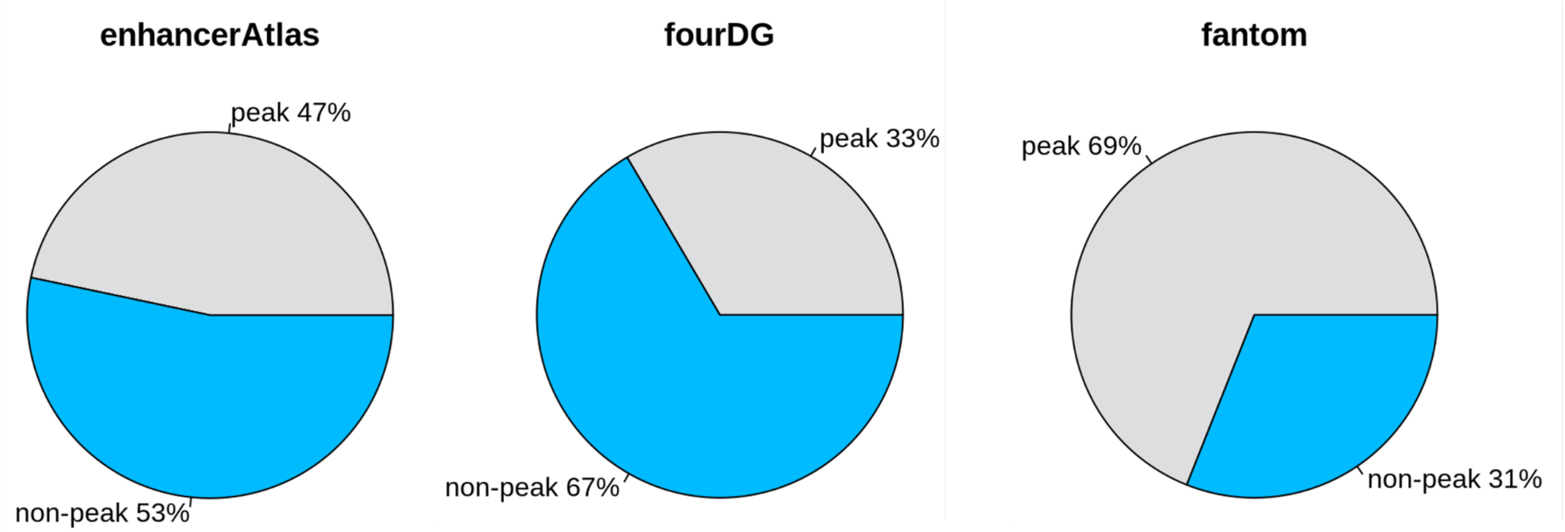

Supp fig 2. Percentage of true regulatory regions recovered by TRIPOD and CREMA when controlling for the precision in the peak regions. Predictions from the two methods were selected at different FDR cutoffs to calculate the precision of regulatory peak prediction and recovery of true regulatory regions from the gold standards.

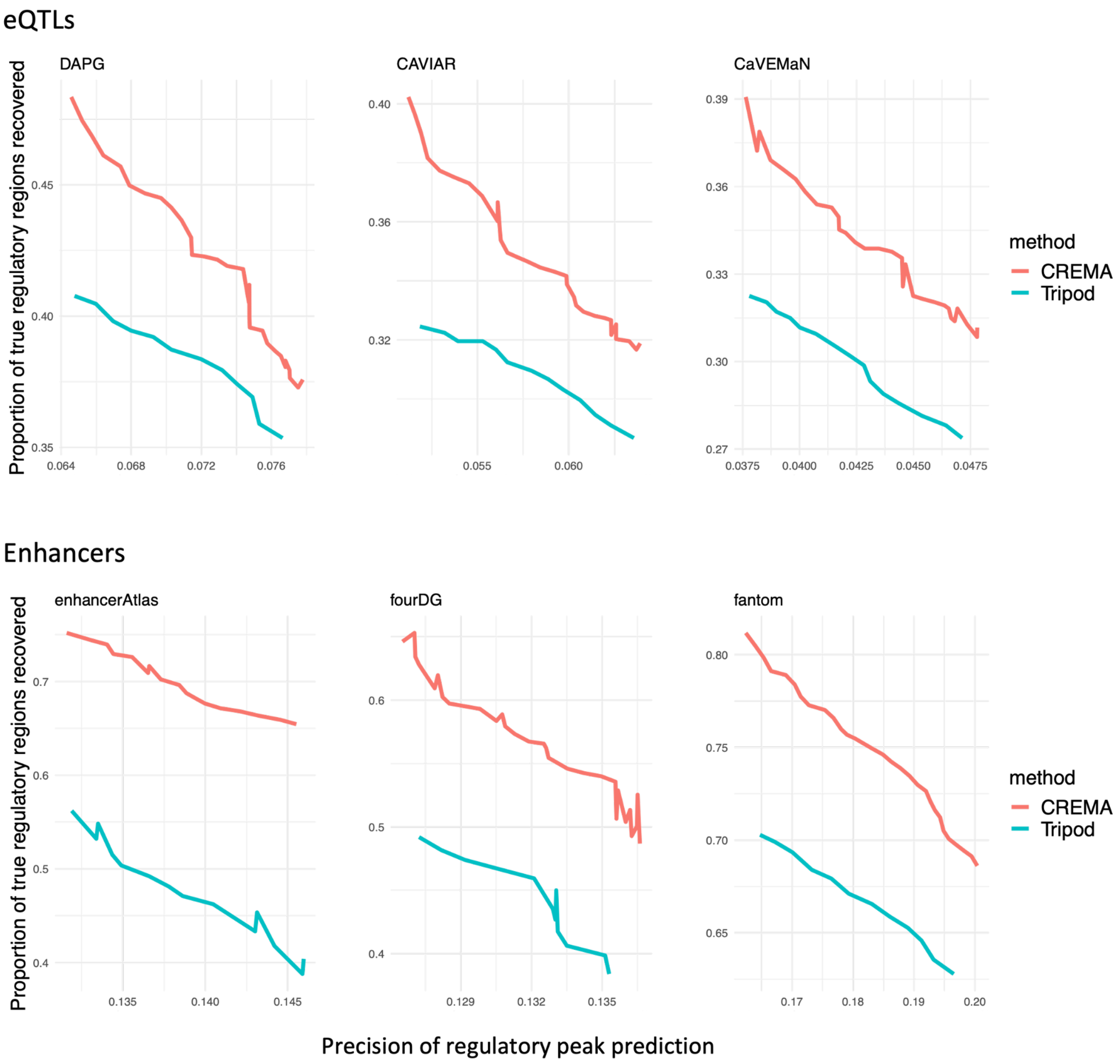

Supplementary figure 3.
Expression of Gata2 in the mouse pituitary tissue (upper) and the corresponding cell type annotations in the same UMAP space (lower).

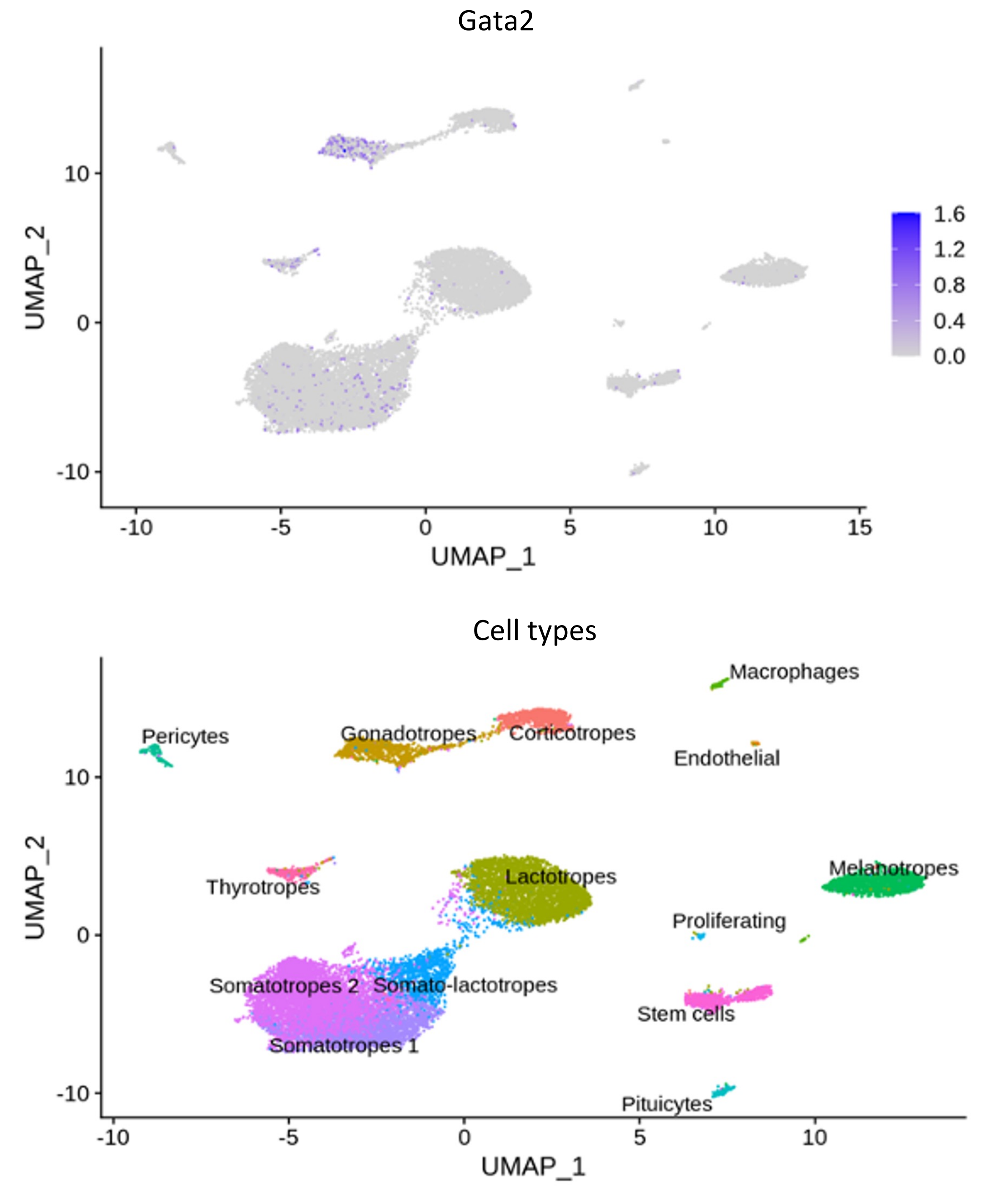

### Supplementary table 1.
### List of TCF7 target genes identified by CREMA in the immune cell types

| celltype | genes |
|---|---|
| Central memory T | CDC14A TNFRSF1B NBPF15 S100A11 FLAD1 ITGB1 ARID5B CD5 ETV6 DUSP16 PLEKHA5 ST8SIA1 KIF21A IFNG-AS1 KLRB1 EPSTI1 LCP1 LPAR6 KLF12 LINC00402 ITGAL MAF ABR RARA CD226 IL4I1 MYO1F DPP4 ICOS EPHA4 WDFY1 FOSL2 GALM EML4 CFAP36 RRBP1 TSHZ2 TMX4 MICAL3 PARVB TIGIT ARHGAP31 MB21D2 KAT2B CMTM8 CMTM6 CCR4 CLDND1 TRIM2 ARAP2 ANTXR2 PTPN13 GPRIN3 ADAM19 SEMA5A MAP3K5 TNFAIP3 ZC3H12D MAP3K4 HLA-DQB1 TBXAS1 NCALD BLK MTSS1 RAB11FIP1 AP3M2 GLIPR2 ANXA1 LINC00892 CD40LG FAAH2 CXCR3 |
| Naive T | SELL ITPKB MDS2 LDLRAP1 MAN1C1 LRRC7 CA6 RGS10 SFMBT2 CRTAM PDE3B PSMA1 PGGHG KLRK1 KLRC4 NELL2 KRT72 NAA16 SLC7A8 ACTN1 LINC01550 APBA2 IGF1R CCR7 NOG PITPNC1 SDK2 CD7 FCGBP ARRDC5 ZNF331 ITGA6 PDK1 FAM117B SNED1 TRABD2A CD8B CD8A SNTG2 PDE9A PRMT2 PIK3IP1 OXNAD1 SATB1 SATB1-AS1 TGFBR2 FHIT FOXP1 LEF1-AS1 LEF1 MAML3 ATP8A1 RAPGEF6 NDFIP1 ARHGAP26 IL6ST RASGRF2 SLC16A10 THEMIS PTPRK CARMIL1 LY86 NT5E BACH2 NRCAM LRRN3 AOAH LEPROTL1 FAM102A MLLT3 |
| Both | NOTCH2 MCL1 CTSS GABPB2 THEM4 RPS27 TPM3 SMG5 USF1 SDHC CD247 CACYBP PTPRC TRAF3IP3 RPL11 NIPAL3 RCAN3 WASF2 LCK RPS8 ZSWIM5 ECHDC2 PATJ RPL5 EVI5 ADD3 USP6NL NMT2 CCDC7 GDI2 PRKCQ PRKCQ-AS1 PSAP SPOCK2 ANAPC16 RPS24 TMEM123 CD3D CD3E CD3G RPS25 ETS1 NAP1L4 CD6 FTH1 FAU MALAT1 PACS1 GSTP1 RPS3 SESN3 BTBD11 RBM19 RPLP0 NCOR2 YARS2 PCED1B-AS1 TESPA1 DGKA RPL41 NACA GNS HELB LYZ SLC2A3 DUSP6 BTG1 LINC01619 CLEC2D TPP2 RASA3 MRPS31 RGCC TPT1 EVL BAZ1A RPS29 KIAA0586 FUT8 SPTLC2 CALM1 CCDC88C TC2N BCL11B TARSL2 B2M MYEF2 USP3 RPLP1 AKAP13 ZNF710 CRTC3 CHD2 TMEM204 RPS15A RPS2 ARHGAP17 LAT CORO7 CYLD RBL2 SLC7A6 WWP2 CYBA RPL13 RPL23A SLFN5 MLLT6 RPL23 SKAP1 ABI3 FAM117A TSPOAP1-AS1 ACAP1 GRB2 RNF157 SEC14L1 TMC8 CHD3 RNF213 VAMP2 CSNK1D LDLRAD4 RNF125 BCL2 TMX3 ARHGAP45 PRDX2 KLF2 RPL18A IFI30 MOB3A KIAA0355 ZNF529 RPS16 EEF2 CEACAM21 POU2F2 FOSB RPL18 FLT3LG NOSIP RPL13A RPS11 AP2A1 ZNF836 RPS9 RPL36 RPS5 CLPP ALKBH7 STXBP2 CFD NCK2 UXS1 GYPC BIN1 CXCR4 ARHGAP15 RBMS1 PRKRA STK17B AC013264.1 IKZF2 LBH ZFP36L2 RHOQ RPS27A CCDC88A CCT4 PPP3R1 AAK1 TET3 MAL ADAM17 ZAP70 MGAT4A SDCBP2-AS1 RBL1 SLC23A2 TMEM230 GPCPD1 CASS4 ZNF831 RPS21 OGFR LINC00649 AGPAT3 TRIOBP RPL3 GRAP2 ST13 SENP7 RPL24 RPL32 TNIK FNDC3B ACAP2 RPL15 SFMBT1 CCDC66 RPL34 ZNF827 FBXL5 DDX60L LAP3 RHOH AFAP1 ST8SIA4 CAMK4 TNFAIP8 CDC42SE2 AFF4 SKP1 IK ANKH RPS14 CD74 ITK CYFIP2 NPM1 SFXN1 MAML1 RACK1 IL7R ANKRD55 ZSWIM6 SERINC5 RPS23 SCML4 STX7 HIVEP2 UTRN ACAT2 IGF2R ERMARD LTA LST1 HLA-DRB5 HLA-DQA1 HLA-DRB1 HLA-DPB1 RPS18 PHF1 SYNGAP1 RPS10 RPL10A STK38 CCND3 EEF1A1 SNHG5 PILRA ZCWPW1 CUX1 NAMPT TMEM106B KDM7A BRAF TRBC2 TRBC1 EPHB6 GIMAP7 STK17A IKZF1 FGL2 GSAP SAMD12 MYC ST3GAL1 EEF1D ASAH1 CHMP7 PTK2B SARAF CEBPD AGPAT5 TNFSF8 RPL35 RPL12 PSIP1 DOCK8 RPL36A RPL39 RPL10 CFP PIM2 PPP1R3F RPS4X |

Supplementary table 2.
table of publicly available data sources used in this study

| Datasets | URL or GEO access number |
|---|---|
| **Publicly available datasets** | |
| Single nucleus multiome of human PBMC | https://www.10xgenomics.com/resources/datasets/pbmc-from-a-healthy-donor-granulocytes-removed-through-cell-sorting-10-k-1-standard-1-0-0 |
| Single nucleus RNAseq and ATACseq of mouse pituitary Gata2 conditional knockout out | GSE190066 |
| **Gold standard databases for evaluation** | |
| DAPG fine-mapped eQTLs (from GTEx v8) | https://storage.googleapis.com/gtex_analysis_v8/single_tissue_qtl_data/GTEx_v8_finemapping_DAPG.tar |
| CAVIAR fine-mapped eQTLs (from GTEx v8) | https://storage.googleapis.com/gtex_analysis_v8/single_tissue_qtl_data/GTEx_v8_fineapping_CAVIAR.tar |
| CaVEMaN fine-mapped eQTLs (from GTEx v8) | https://storage.googleapis.com/gtex_analysis_v8/single_tissue_qtl_data/GTEx_v8_finemapping_CaVEMaN.tar |
| EnhancerAtlas v2.0 | http://www.enhanceratlas.org/downloadv2.php |
| 4D Genome | https://github.com/yuchaojiang/TRIPOD |
| Fantom v5 | https://github.com/yuchaojiang/TRIPOD |
| **New datasets** | |
| Mouse pituitary single nucleus multiome | GSE234943 |
